# IL-2 enhances effector function but suppresses follicular localization of CD8^+^ T cells in chronic infection

**DOI:** 10.1101/2024.05.02.592184

**Authors:** Yaping Chen, Pengcheng Zhou, Patrick Marc Gubser, Yew Ann Leong, Jing He, Yunbo Wei, Fadzai Victor Makota, Mehrdad Pazhouhandeh, Ting Zheng, Joseph Yunis, Zhanguo Li, Axel Kallies, Di Yu

## Abstract

Cytotoxic CD8^+^ T cells, essential in combating viral infections and cancer, become dysfunctional from prolonged antigen exposure. Precursors of exhausted T (T_PEX_) cells are pivotal in sustaining immune responses in chronic diseases and mediating immunotherapy efficacy. They also control viral infection within B-cell follicles, facilitated by CXCR5 expression. How cytokines regulate T_PEX_ cell fate and follicular entry is not well understood. We reveal that IL-2 treatment enhances CD8^+^ T cell effector functions in chronic LCMV infection but hinders CXCR5^+^ T_PEX_ cell formation and infection control within B-cell follicles. Mechanistically, IL-2 suppresses T_PEX_ cell differentiation in a STAT5 and BLIMP1-dependent manner. Using an IL-2 fusion protein targeting CD122, we shifted the differentiation towards CX3CR1^+^ T cells with increased effector function. Clinical observations with low-dose IL-2 in autoimmune disease confirmed IL-2’s inhibitory effect on CXCR5^+^ T_PEX_ cells, underscoring IL-2’s crucial regulatory role and therapeutic potential in modulating T_PEX_ and effector T cell generation.

## INTRODUCTION

CD8^+^ T cells play a critical role in eliminating virally infected and malignant cells. In chronic infection and cancer, antigen-specific CD8^+^ T cells persistently exposed to antigen gradually reduce their ability to produce effector cytokines and proliferative capacity. This process is paralleled by the upregulation of inhibitory receptors or checkpoints such as programmed cell death protein 1 (PD-1), T cell immunoglobulin and mucin domain 3 (TIM-3) and lymphocyte-activation gene 3 (LAG3). Although such cellular reprogramming is necessary to restrain CD8^+^ T cells-mediated immunopathology, reduced functionality in CD8^+^ T cells impairs the clearance of infection or prevention of tumor growth. This process is referred to as T cell exhaustion (Blank et al., 2019; Hashimoto et al., 2018; McLane et al., 2019).

PD-1^+^ exhausted CD8^+^ T cells are heterogenous and contain a population of cells known as progenitors or precursors of exhausted T (T_PEX_) cells. These cells retain the capacity to self-renew while also giving rise to exhausted effector T (T_EX_) cells as we and others have shown (He et al., 2016b; Im et al., 2016; Leong et al., 2016; Utzschneider et al., 2016; Wu et al., 2016). T_PEX_ cells also mediate the response to immune checkpoint blockade (ICB) therapies that inhibit PD-1/PD-Ligand 1 (PD-L1) interactions and thus amplify anti-viral and anti-tumor immunity in mice and humans (Eberhardt et al., 2021; Huang et al., 2022; Im et al., 2016; Siddiqui et al., 2019; Utzschneider et al., 2016). T_PEX_ cells express the chemokine receptor CXCR5, which facilitates their migration to B-cell follicles where they contribute to the suppression of infection (He et al., 2016b; Leong et al., 2016). This is important as B-cell follicles act as reservoirs for certain viruses, such as Epstein–Barr virus (EBV) in memory B cells and Human Immunodeficiency Virus (HIV) in follicular helper T (T_FH_) cells (Collins et al., 2023; Leong et al., 2016; Mylvaganam et al., 2017; Petrovas et al., 2017). Therefore, CXCR5-expressing T_PEX_ cells are critical in suppressing the infection in B-cell follicles (Yu and Ye, 2018).

T cell exhaustion is controlled by a network of transcriptional regulators induced by T cell receptor (TCR) stimulation-induced transcription factors, in particular NFATC1, TOX and IRF4 drive the sustained expression of inhibitory receptors and contribute to the downregulation of effector function (Alfei et al., 2019; Khan et al., 2019; Man et al., 2017; Martinez et al., 2015; Scott et al., 2019; Seo et al., 2019; Yao et al., 2019). The generation and maintenance of T_PEX_ cells, on the other hand, are specifically controlled by transcription factors such as TCF1, MYB, ID3 and BCL6 while T_EX_ cells differentiation is promoted by BLIMP1(Gautam et al., 2019; He et al., 2016b; Im et al., 2016; Leong et al., 2016; Tsui et al., 2022; Utzschneider et al., 2016; Wu et al., 2016). However, extracellular signals that regulate T_PEX_ and T_EX_ cell differentiation and maintenance are less understood. We and others have shown that TGFβ plays a key role in maintaining T_PEX_ cell function and identity (Gabriel et al., 2021; Hu et al., 2022; Ma et al., 2022). Furthermore, both T_FH_ cells-derived cytokine interleukin-21 (IL-21) and the alarmin IL-33 are necessary to maintain T_PEX_ cells and sustain effector CD8^+^ T cell function in chronic infection (Marx et al., 2023; Yu et al., 2022; Zander et al., 2022). Finally, IL-2, in combination with ICB, was shown to direct T_PEX_ cells to generate transitory effector CD8^+^T cells and improve antiviral and antitumor activities (Codarri Deak et al., 2022; Hashimoto et al., 2022; Ren et al., 2022; Tichet et al., 2023; West et al., 2013). However, the precise function of IL-2 in regulating T cell exhaustion is not fully understood.

To better understand how IL-2 regulates T_PEX_ cell differentiation and function, we investigated the impact and molecular mechanisms of treatment with IL-2 or a clinical-grade IL-2 fusion protein specifically targeting the IL-2 receptor β chain (CD122) in chronic LCMV infection. IL-2 treatment increased the generation of CX3CR1+ T_EX_ cells with enhanced effector function without disrupting the self-renewal capacity of T_PEX_ cells both in mouse and human. At the same time, IL-2 treatment resulted in downregulation of CXCR5 expression on T_PEX_ cells, their exclusion from B-cell follicles and impaired viral clearance from these sites. Furthermore, increased IL-2 signaling during the early stages of the immune response strongly suppressed T_PEX_ cell generation. Mechanistically, IL-2 exerted its effects in a STAT5– and BLIMP1-dependent manner. Overall, our data show that IL-2 treatment can broadly modulate the balance between effector T cells and T_PEX_ cells and excert differential effects on viral clearance.

## RESULTS

### IL-2 treatment promotes CD8^+^ T effector expansion during chronic infection

IL-2 is an approved immunotherapeutic agent for cancer (Dutcher et al., 2014). It’s precise mechanism of action is not fully understood. To elucidate how IL-2 affects exhausted CD8^+^ T cell maintenance and function, wild-type C57BL/6 mice were infected with LCMV Docile (10^6^ plaque-forming units, PFU), which induces chronic infection and robust T cell exhaustion. Mice were treated with PBS or low-to-medium dose of recombinant human IL-2 (30,000 international units, IU (Blattman et al., 2003; He et al., 2016a; West et al., 2013)) daily from day 10 to 14 post-infection before analysis on day 15 (**Figure 1A**). IL-2 treatment elevated CD8^+^ T cell numbers and increased CD8^+^/CD4^+^ T cell ratios (**Figure 1B**). Furthermore, numbers of activated CD8^+^ T cells (CD44^high^CD62L^-^) and their effector function as measured by Granzyme B expression were significantly enhanced by IL-2 treatment (**Figure 1C, D**). In agreement with previous studies (Blattman et al., 2003; Zhou et al., 2021), we found that IL-2 treatment significantly increased both the frequencies and numbers of LCMV-specific CD8^+^ T cells as measured by staining with H-2D^b^ LCMV GP_33_ tetramers (**Figure 1E**). We did not observe an expansion of regulatory (T_REG_) cells by this dose of IL-2 treatment (data not shown), presumably due to the high numbers of activated effector T cells (Humblet-Baron et al., 2016; Zhou et al., 2021). We next examined the impact of IL-2 treatment on the generation of TCF1^+^CXCR5^+^ T_PEX_ and TCF1^-^CXCR5^-^ T_EX_ cells. IL-2 treatment increased the total numbers of T_EX_ cells by more than two-fold (**Figure 1F**). In contrast, IL-2 treatment reduced the frequencies of T_PEX_ cells within GP_33_ tetramer^+^ CD8^+^ T cells by more than two-fold although their numbers remained largely unchanged (**Figure 1F**). Overall, IL-2 treatment enhanced effector T cell expansion in chronic infection.

**Figure 1.**
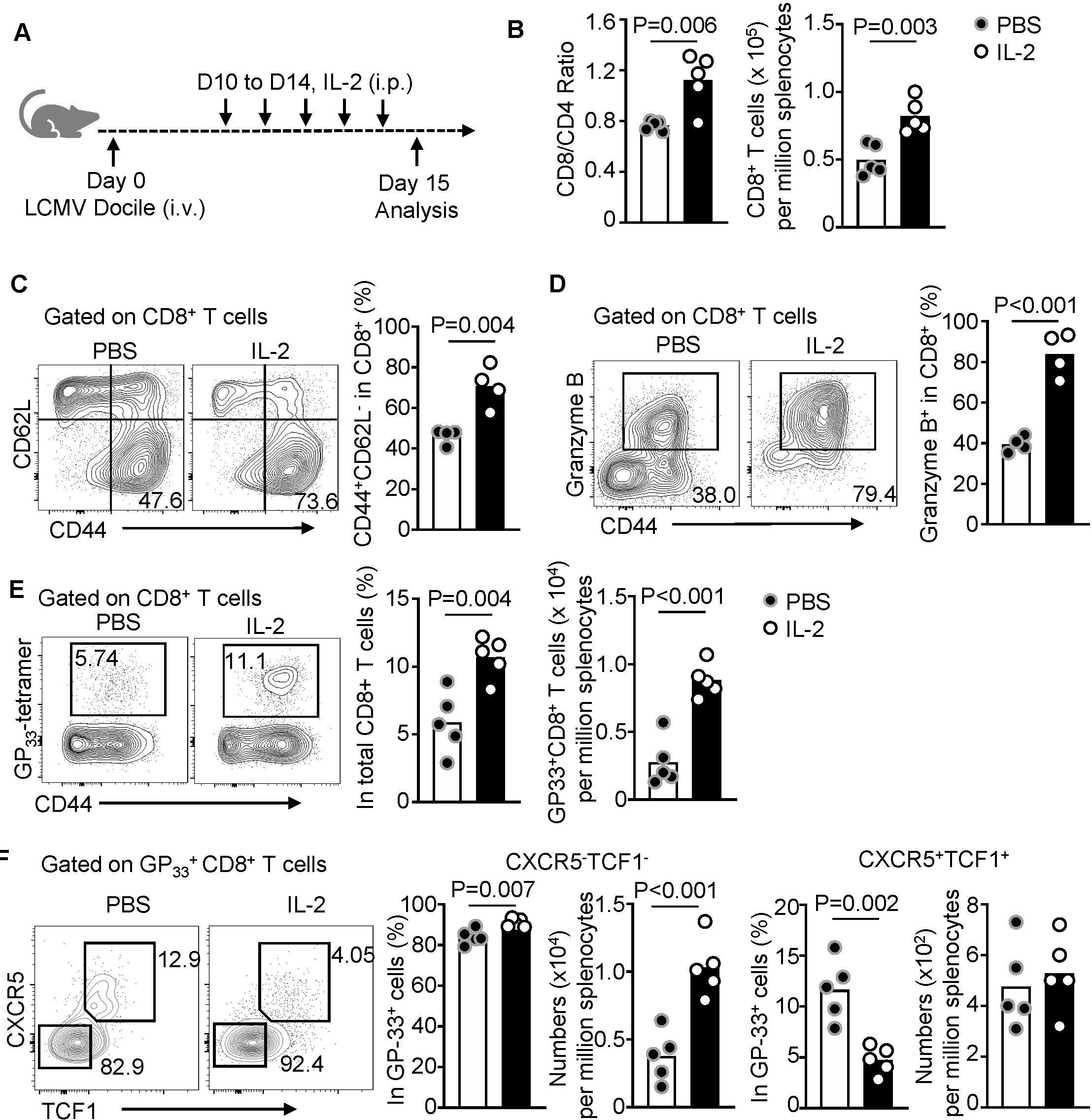
IL-2 treatment promotes CD8+ T effector expansion during chronic infection. (**A**) Schematic showing C57BL/6 mice chronically infected with LCMV Docile and receiving intraperitoneal treatment of 30,000 IU recombinant human IL-2 daily from day 10 to 14 post infection, followed by the analyses on day 15. (**B**) Quantification showing CD4/CD8 ratio and the numbers of CD8^+^ T cells in the spleen. Cells were gated on 7AAD^-^ B220^-^ TCRβ^+^CD8^+^ or CD4^+^ cells using flow cytometry. (**C**,**D**) Representative FACS plots and statistics showing CD44^high^CD62L^low^ effector (C) and the expression of Granzyme B (D) in total CD8^+^ T cells in mice treated with PBS or IL-2. (**E, F**) Representative FACS plots and quantification showing the frequencies and numbers of H-2Db LCMV GP_33_ tetramer-specific CD8^+^ T cells (E) and the frequencies and numbers of CXCR5^-^TCF1^-^ T_EX_ cells and CXCR5^+^TCF1^+^ T_PEX_ cells. Results are representative of two or three independent experiments (*n* = 4-5/group). Each dot represents one individual mouse. Student’s unpaired two-tailed t tests.

### IL-2 alters tissue localization of CD8^+^ T cells and viral reservoirs in chronic infection

Previous studies reported that a short period of IL-2 treatment, despite enhancing CD8^+^ T effector function, was insufficient to significantly reduce viral loads (Blattman et al., 2003; Zhou et al., 2021). LCMV preferentially infects myeloid cells expressing high levels of α-Dystroglycan (α-DG), the cellular receptor for LCMV (Kunz et al., 2001; Oldstone and Campbell, 2011), but it also infects lymphocytes including CD4^+^ T cells (Leong et al., 2016). Immunofluorescent microscopy of splenic tissue using the antibody (VL4) to LCMV nucleoprotein revealed that IL-2 treatment resulted in an altered distribution of VL4^+^ LCMV reservoirs towards B220^+^ B-cell follicles (**Figure 2A**). To examine whether IL-2 treatment influenced the distribution of virus in the B-cell follicle and the T-cell zone, we deployed a flow cytometry-based method previously developed (Leong et al., 2016). Treatment with IL-2, in a dose-dependent manner, reduced numbers of virus-infected CXCR5^low^CD44^+^ non-T_FH_ effector CD4^+^ T cells, which primarily reside in the T-cell zone, whereas the numbers of virally infected CXCR5^high^CD44^+^ T_FH_ cells, which mostly reside in B-cell follicles, remained unchanged. As a consequence, the LCMV reservoir was biased towards the B-cell follicle, indicated by a two-fold increase in the ratio of infected T_FH_/non-T_FH_ cells at the highest dose of IL-2 treatment (**Figure 2B**).

**Figure 2.**
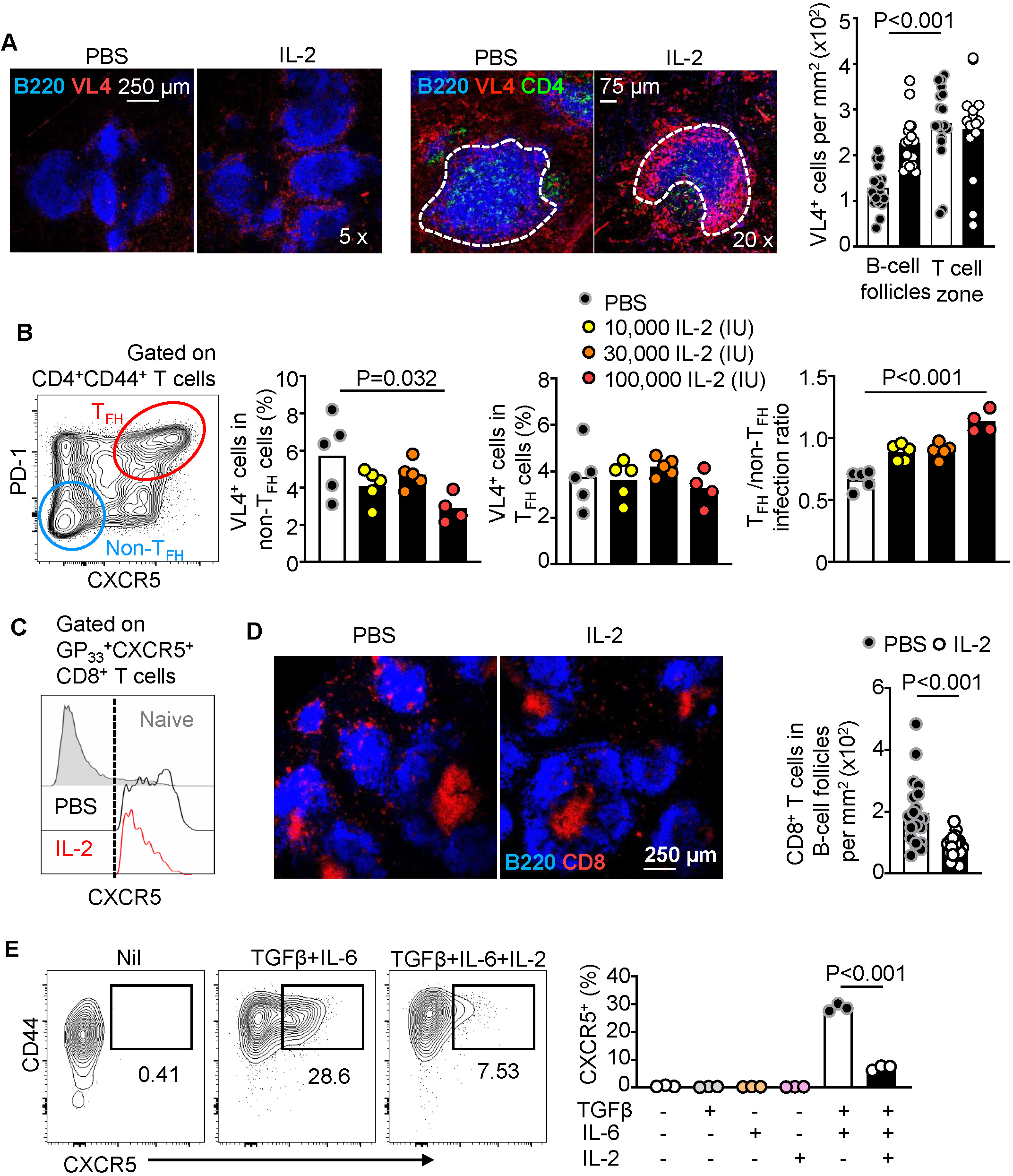
IL-2 alters tissue localization of CD8^+^ T cells and viral reservoirs in chronic infection. (**A**) C57BL/6 mice were chronically infected with LCMV Docile and received intraperitoneal treatment with IL-2 as in Figure 1A. Representative confocal images (left: low magnification, right: high magnification) and quantification of LCMV infected cells in the spleen. (**B**) C57BL/6 mice chronically infected with LCMV Docile and receiving intraperitoneal treatment of gradient doses of recombinant human IL-2 (10,000 – 100,000 IU) daily from day 10 to 14 post infection, followed by the analyses on day 15. FACS plot showing the gating of T_FH_ and non-T_FH_ cells in CD4^+^ T cells (left), and quantification of virus-infected (VL4^+^) T_FH_ and non-T_FH_ cells in each group (right). (**C**, **D**) In the same experiment as (A), histograms showing the expression of CXCR5 on GP_33_ tetramer-specific CXCR5^+^ CD8^+^ T cells (C) and representative images (left) and quantification (right) of CD8^+^ T cells in B-cell follicles (D). (**E**) Representative FACS plots (left) and quantification (right) showing the expression of CXCR5^+^ on OT-I CD8^+^ T cells, co-cultured with splenocytes pulsed with the SINFEKL peptide (1 µM) and indicated cytokines: TGFβ (10 ng/mL), IL-6 (100 ng/mL) and IL-2 (500 I.U./mL). Results are representative of at least two independent experiments. Each dot represents one image count (A, D), one individual mouse (B), or one culture (E). Student’s unpaired two-tailed t tests.

T_PEX_ cells that express CXCR5 have access to the B-cell follicle and contribute to viral control (Collins et al., 2023; Leong et al., 2016; Petrovas et al., 2017). The altered distribution of LCMV reservoirs, however, suggested that IL-2 treatment prevents access of CD8^+^ T cells to B-cell follicles. Indeed, IL-2 treatment significantly reduced CXCR5 expression on T_PEX_ cells (**Figure 2C**). In line with this observation, immunofluorescent imaging of splenic tissue sections revealed a drastic reduction of CD8^+^ T cells in the B-cell follicles in IL-2-treated mice compared with that of PBS-treated control mice (**Figure 2D**).

Systemic administration of IL-2 may control CD8^+^ T cell activation and differentiation directly or indirectly. We therefore assessed the direct impact of IL-2 *in vitro*. Stimulation of human CD8^+^ T cells with anti-CD3/CD28 and cytokines TGFβ, IL-12, and IL-23 was shown to promote CXCR5 expression (Mylvaganam et al., 2017). In lijne with that observation, we and others found that TGFβ supported T_PEX_ cell generation and maintenance (Gabriel et al., 2021; Hu et al., 2022; Ma et al., 2022). We therefore optimized a culture condition whereby naïve mouse OT-I CD8^+^ T cells that carry a transgenic TCR-specific to ovalbumin (OVA) antigen were stimulated with splenocytes pulsed with SIINFEKL peptide in the presence or absence of TGFβ and IL-6, with or without IL-2. The combination of TGFβ and IL-6 induced CXCR5 expression on OT-I T cells. In contrast, the addition of IL-2 in the culture diminished CXCR5 expression by about three-fold (**Figure 2E**). Overall, these results suggest that IL-2 induces signaling that favors the generation of CXCR5^-^ T_EX_ cells over CXCR5^+^ T_PEX_ cells in CD8 T cells.

### An IL-2-STAT5-BLIMP1 axis regulates the balance of effector and T_PEX_ generation

To better understand the impact of IL-2 on T_PEX_ and T_EX_ cell generation, we performed Gene Set Enrichment Analysis (GSEA) of CXCR5^+^ T_PEX_ cells compared to CXCR5^-^ T_EX_ cells (Leong et al., 2016). This analysis indicated downregulation of IL-2-STAT5 signaling in CXCR5^+^ T_PEX_ cells (**Suppl. Figure 1A**), suggesting that IL-2 negatively impacts the generation or function of CXCR5^+^ T_PEX_ cells. The IL-2 receptor is a heterotrimeric protein complex composed of three subunits: the α subunit CD25 encoded by *Il2ra*, the β subunit CD122 encoded by *Il2rb*, and the common γ chain CD132 encoded by *Il2rg* (Boyman and Sprent, 2012). Transcriptomic analysis indicated that *Il2ra* and *Il2rb* were lower in CXCR5^+^ T_PEX_ population, whereas *Il7r* was higher, compared with CXCR5^-^ T_EX_ cells (**Suppl. Figure 1B,C**). Flow cytometric analysis showed that the expression of CD122 on CXCR5^+^ T_PEX_ was significantly lower than that on CXCR5^-^ T_EX_ cells, while CD25 protein was minimally expressed on both populations (**Figure 3A**). Notably, the treatment of IL-2 in mice with LCMV infection pronouncedly increased CD122 expression and modestly increased CD25 expression on T_PEX_ cells, whereas such effects were not observed in T_EX_ cells (**Figure 3B** and **Suppl. Figure 1D**).

**Figure 3.**
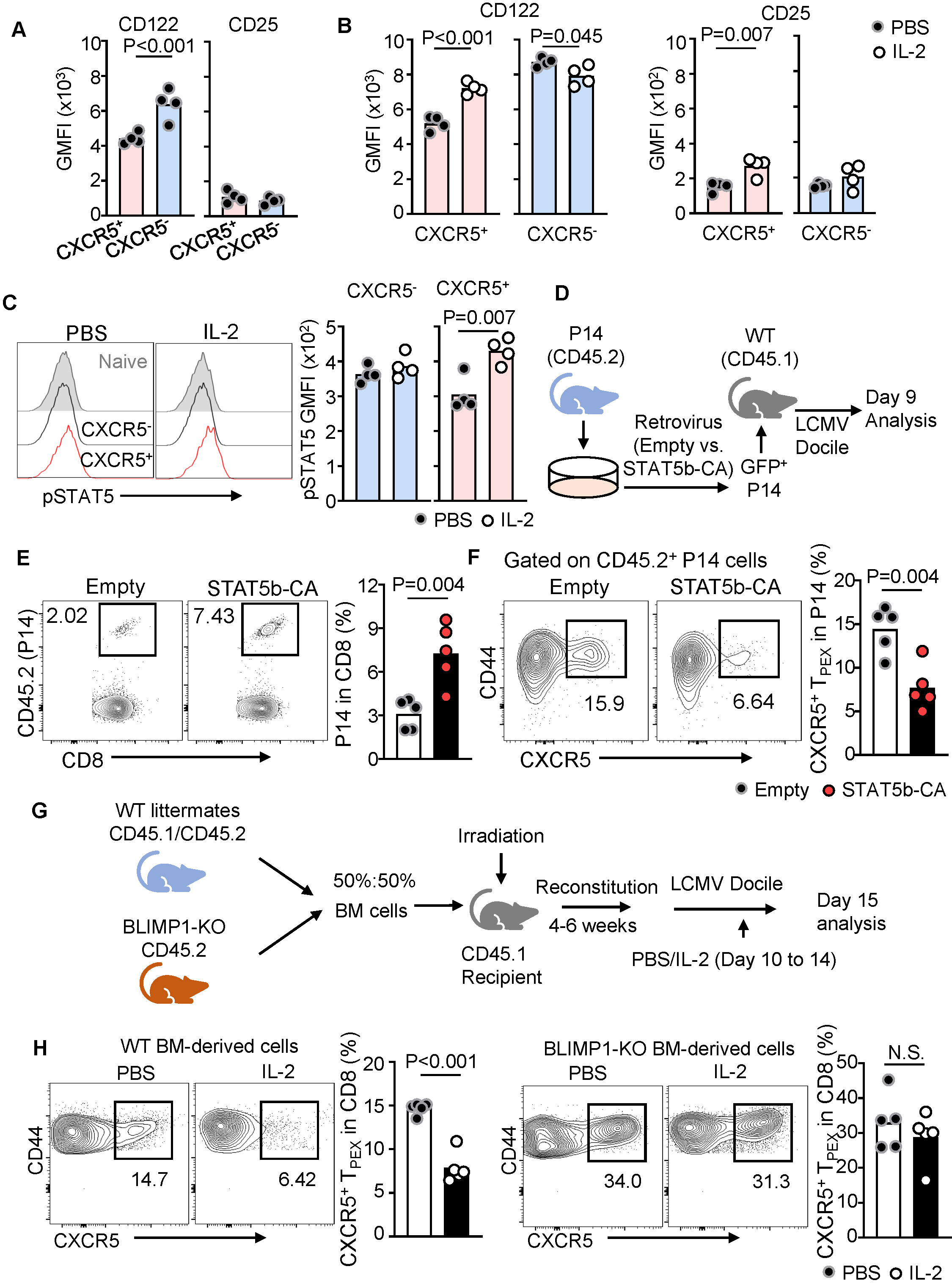
An IL-2-STAT5-BLIMP1 axis regulates the balance of effector and T_PEX_ generation. (**A-C**) C57BL/6 mice were chronically infected with LCMV Docile and received intraperitoneal treatment with IL-2 as in Figure 1A. Quantification of the expression of CD122 and CD25 between CXCR5^+^ T_PEX_ cells and CXCR5^-^ T_EX_ cells (A). The comparison of the expression of CD122 and CD25 on CXCR5^+^ T_PEX_ cells or CXCR5^-^ T_EX_ cells between PBS treated and IL-2 treated mice (B). The comparison of the expression of pSTAT5 in CXCR5^+^ T_PEX_ cells or CXCR5^-^ T_EX_ cells between PBS treated and IL-2 treated mice (C). (**D-F**) Schematic (D) showing the overexpression of constitutive active STAT5b (STAT5b-CA) in P14 cells, followed by the adoptive transfer in congenically marked recipient mice with chronical LCMV Docile infection. Representative FACS plots (left) and quantifications (right) of the frequencies of P14 cells in total CD8^+^ T cells (E) and the frequencies of CXCR5^+^ T_PEX_ cells in P14 cells (F). (**G**, **H**) Schematic of the generation of bone marrow chimera mice carrying T cell specific deletion of BLIMP1 (encoded by *Prdm1*), followed with chronic LCMV Docile infection and IL-2 treatment (i.p. 30,000 I.U. daily from day 10 to 14 post-infection) (G). Representative FACS plots and quantifications showing the frequencies of CXCR5^+^ T_PEX_ cells in CD44^+^ CD8^+^ T cells (H). Results are representative of two or three independent experiments (*n* = 4-5/group). Each dot represents one individual mouse. Student’s unpaired two-tailed t tests. N.S., statistically not significant.

The phosphorylation of STAT5 (pSTAT5) is a major signaling event induced by IL-2 (Boyman and Sprent, 2012). IL-2 treatment increased the pSTAT5 level in CXCR5^+^ T_PEX_ cells but not in CXCR5^-^ T_PEX_ cells (**Figure 3C**). To directly test the impact of STAT5 signaling, we transduced CD45.2 TCR transgenic P14 T cells that are specific for the LCMV GP_33-41_ epitope with the retrovirus expressing a constitutively activated version of STAT5b (STAT5b-CA) or an empty viral vector control (Johnston et al., 2012). Transduced P14 T cells were then transferred into congenically marked CD45.1 recipient mice followed by LCMV Docile infection (**Figure 3D**). Similar to the effects of IL-2 treatment and in line with a recent report (Beltra et al Immunity 2023), STAT5b-CA overexpression led to an increase in total P14 cells by about three-fold compared to controls (**Figure 3E**). Importantly, the constitutive activation of the STAT5 signaling in P14 cells favored the generation of CXCR5^-^ T_EX_ over CXCR5^+^ T_PEX_ cells *in vivo* (**Figure 3F**).

CXCR5^+^ T_FH_ cells and CXCR5^+^ T_PEX_ cells share a similar transcriptional program for their generation whereby both are promoted by transcription factors TCF1 and BCL6 and suppressed by BLIMP1 and ID2 (Kallies et al., 2020; Vinuesa et al., 2016). We and others previously reported that IL-2/STAT5 signaling induces the expression of B lymphocyte-induced maturation protein-1 (BLIMP1, encoded by *Prdm1*), a negative regulator of *Cxcr5* transcription (Johnston et al., 2009; Leong et al., 2016; Xin et al., 2016). We therefore examined the roles of BLIMP1 in regulating the differentiation of CXCR5^+^ T_PEX_ cells in response to IL-2. First, we generated mixed bone marrow chimeric mice containing cells from mice carrying a T cell-specific ablation of Blimp1 (*Prdm1*^flox/flox^*Lck*^Cre^) and cells from congenically marked wildtype mice, mixed at a 1:1 ratio. These chimeras were infected with LCMV Docile, treated with IL-2 or PBS from day 10 to day 14 and analyzed on day 15 (**Figure 3G**). As expected, we observed larger proportions of CXCR5^+^ T_PEX_ cells in BLIMP1-deficient (∼30%) compared to control (∼15%) CD8^+^ T cells. Notably, IL-2 treatment further reduced the CXCR5^+^ T_PEX_ cell percentages from ∼15% to ∼6% in control CD8^+^ T cells but did not reduce T_PEX_ cell frequencies in BLIMP1-deficient CD8^+^ T cells, which remained as ∼30% (**Figure 3H**). Taken together, our results indicate that the IL-2-STAT5-BLIMP1 axis plays a critical role in regulating the generation of CXCR5^+^ T_PEX_ cells in response to IL-2 treatment in chronic infection.

### An IL-2 fusion protein regulates the balance of effector and T_PEX_ cell generation

IL-2 fusion proteins that increase the half-life or bioavailability of IL-2 have recently been shown to be promising tools in cancer immunotherapy (Beltra et al., 2023; Codarri Deak et al., 2022; Hashimoto et al., 2022; Tichet et al., 2023). To specifically assess the role of the IL-2-CD122 axis in regulating the generation of T_PEX_ cells, we examined the effects of a novel IL-2 fusion protein, ANV410, that selectively targets CD122. Consistent with the CD122-specific activity, treatment of naïve mice with ANV410 resulted in a dramatic expansion of CD44^+^ memory CD8^+^ T cells, and NK cells, without affecting T_REG_ cells (**Suppl. Figure 2**). We next treated mice chronically infected with LCMV Docile (8-12 days post infection), which had received LCMV-specific P14 T cells (Regimen A, **Figure 4A**). Total CD8^+^ as well as P14 T cells expanded substantially in response to ANV410 (**Figure 4B**). We also observed a significant increase in the percentage and numbers of CX3CR1^+^ T_EX_ cells (**Figure 4C**), which harbor the highest functional capacity among exhausted T cells (Hudson et al., 2019; Zander et al., 2019). Consistent with this observation, we also found larger numbers of P14 T cells expressing Granzyme B (**Figure 4D**). In contrast, the percentages of T_PEX_ cells in P14 T cells declined while their numbers remained unchanged (**Figure 4E**), which was consistent with the results from IL-2 treatment (**Figure 2E**). To examine the effects of the IL-2 fusion protein on the generation of T_PEX_ cells, we treated mice with ANV410 during the first 5 days post infection (Regimen B, **Figure 4A**). Strikingly, early ANV410 treatment dramatically impaired the expansion of P14 T cells (**Figure 4F**) and abrogated TPEX cell development almost completely both in the P14 and polyclonal PD-1^+^ CD8 T cell compartment (**Figure 4G, H**), suggesting that increased IL-2 signaling early during infection is detrimental to T_PEX_ cell development and function.

**Figure 4.**
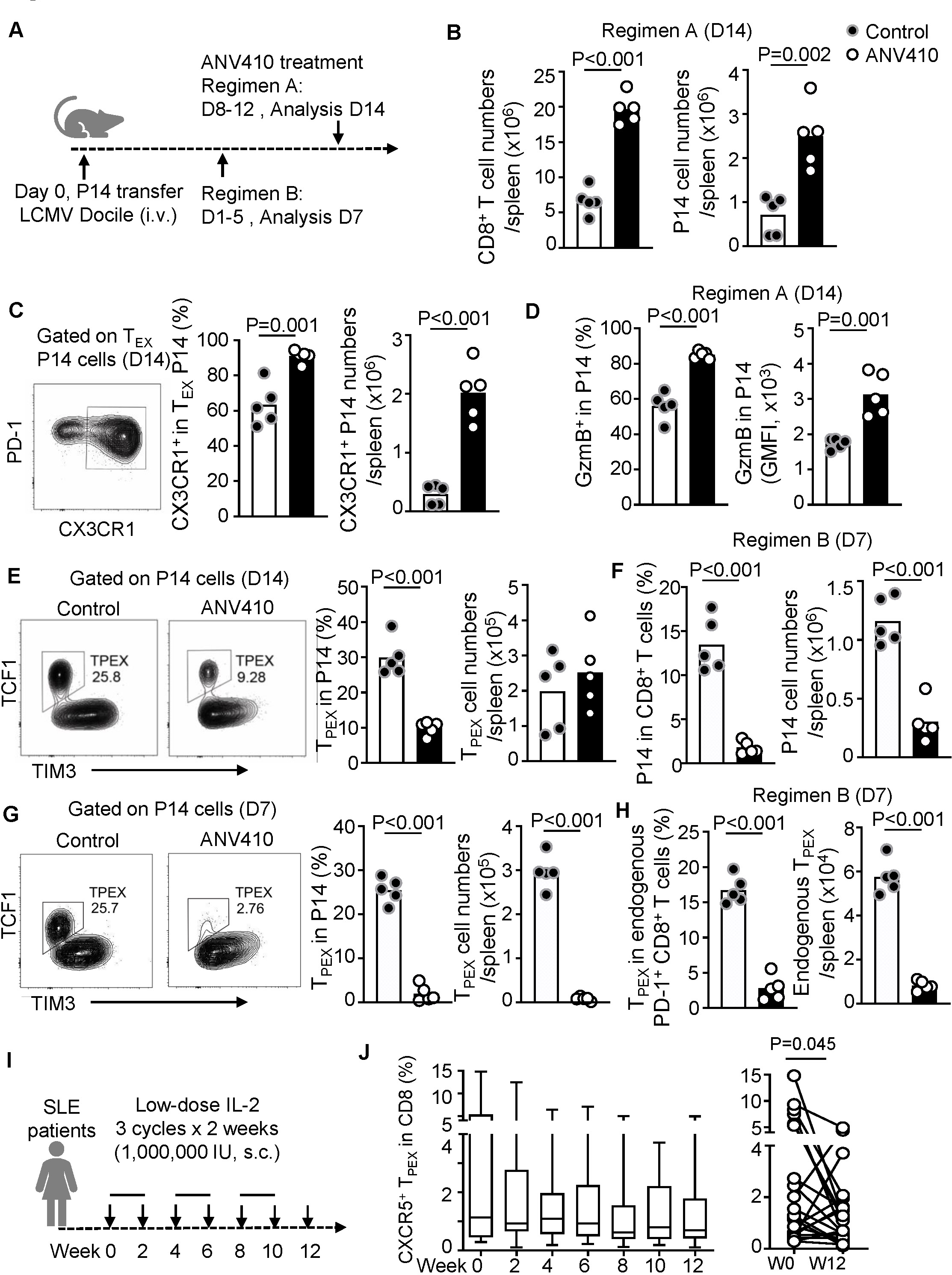
An IL-2 fusion protein regulates the balance of effector and T_PEX_ cell generation. (**A**) Schematic showing P14 T cells being transferred into congenically marked mice, which were subsequently infected with LCMV Docile and treated with ANV410, a CD122-specific IL-2 fusion protein, three times every other day from day 8 to 12 post infection and analyzed on day 14 post infection (regimen 1, panels B-E) or from day 1 to 5 of infection and analyzed on day 7 post infection (regimen 2, panels F-H). (**B**) Quantifications of total CD8^+^ T cells (left) or P14 cells (right) on day 14 post infection (regimen 1). (**C**) FACS plot showing the gating strategy for CX3CR1^+^ T_EX_ cells (left) and quantifications of percentages and numbers of CX3CR1^+^ T_EX_ cells in P14 cells on day 14 post infection (regimen 1). (**D**) Quantification showing percentages and expression levels of GzmB^+^ P14 cells on day 14 post infection (regimen 1). (**E**) FACS plots showing the gating strategy for TCF1^+^ T_PEX_ cells (left) and quantification showing the percentages and numbers of TCF1^+^ T_PEX_ cells in P14 cells on day 14 post infection (regimen 1). (**F**) Quantification showing the percentages and numbers of P14 cells on day 7 post infection (regimen 2). (**G**) FACS plots showing the gating strategy for TCF1^+^ T_PEX_ cells (left) and quantification showing the percentages and numbers of TCF1^+^ T_PEX_ cells in P14 cells on day 7 post infection (regimen 2). (**H**) Quantification showing the percentages and numbers of TCF1^+^ T_PEX_ cells in endogenous PD-1^+^ CD8^+^ T cells on day 7 post infection (regimen 2). (**I**) Schematic of low-dose IL-2 therapy in patients with systemic lupus erythematosus (SLE). (**J**) Quantification showing the frequencies of CXCR5^+^ T_PEX_ in peripheral blood CD8^+^ T cells in consecutive time points (left) or before and after the therapy (right). Each dot represents one individual mouse (B-H) or one individual patient (J). Student’s unpaired two-tailed t tests (B-H) and paired two-tailed t test (J).

### IL-2 treatment reduces CXCR5^+^ T_PEX_ cells in humans

Finally, we asked whether IL-2 treatment in humans could regulate T_PEX_ cells. We took advantage of immunological profiling data that we previously collected in a clinical trial of low-dose IL-2 therapy in patients with systemic lupus erythematosus (SLE). In this clinical trial, patients were treated with three two-week cycles of 1,000,000 IU recombinant human IL-2 every other day (**Figure 4I**) (He et al., 2016a). The results demonstrated that low-dose IL-2 treatment significantly reduced human CXCR5^+^ T_PEX_ cells in the blood after three cycles of low-dose IL-2 therapy (**Figure 4J**); this is in agreement with the observation of suppression of CXCR5 expression on CD8^+^ T cells by IL-2 treatment in the mouse model of LCMV chronic infection (**Figure 2C**). Overall, our data in both mice and humans indicate that increased IL-2 signaling via CD122 during early stages of the infection can impair the generation of T_PEX_ cells, while a short period of therapeutic IL-2 treatment at later stages of the infection increases the differentiation of effector cells from T_PEX_ cells without impacting their self-renewal capacity.

## DISCUSSION

Upon chronic antigen exposure, CD8^+^ T cells upregulate the expression of inhibitory receptors including PD-1. PD-1^+^ T cells in chronic infection are heterogenous, containing TCF1^+^ T_PEX_ cells that undergo self-renewal but also give rise to TCF1^-^ T_EX_ cells that lack long-term proliferative capacity (He et al., 2016b; Huang et al., 2022; Im et al., 2016; Leong et al., 2016; Utzschneider et al., 2016; Wu et al., 2016). Critically, T_PEX_ cells mediate the proliferative burst in response to PD-1 immune checkpoint inhibition, which induces the differentiation of CX3CR1^+^ effector-like T_EX_ cells, which display the highest effector function and antiviral activity and closely correlate with positive response to therapeutic ICB in cancer (Hudson et al., 2019; Yan et al., 2018; Zander et al., 2019). Maintaining a balance between self-renewal, effector function and terminal exhaustion underlying by these functional subsets is essential to sustain cytotoxic killing while restraining immunopathology at the same time (Blank et al., 2019; Hashimoto et al., 2018; Kallies et al., 2020; McLane et al., 2019). Our results, together with studies published recently (Beltra et al., 2023; Codarri Deak et al., 2022; Hashimoto et al., 2022; Liu et al., 2021; Ren et al., 2022; Sun et al., 2023) suggest that regulating the T_EX_ and T_PEX_ cell balance is a central role of IL-2 and the downstream STAT5-BLIMP1 axis. Importantly, IL-2 promoted the generation of CX3CR1^+^ effector-like T_EX_ cells without compromising the self-renewal capacity of T_PEX_ cells.

IL-2 signaling is mediated by both the high affinity receptor IL-2Rα (CD25) and the medium affinity receptor IL-2Rβ (CD122). Their individual contribution, however, is dependent on their respective expression levels as well as the dose of IL-2. We found that treatment with native IL-2 resulted in the upregulation of CD122 and induced feedforward signaling. Consistent with this observation, the IL-2 fusion protein ANV410 that selectively targets CD122 but not CD25 induced the generation of CX3CR1 effector-like T_EX_ cells, overall suggesting that CD122 is the main mediator of IL-2 effects on T_PEX_ cells. These results align well with recent studies, which showed that CD122-dependent signals drive terminal differentiation of CD8^+^ T cells (Beltra et al., 2016) and CD122-biased IL-2 agonism in the combination with ICB promoted CD8^+^ T cell effector function (Codarri Deak et al., 2022). Our results, however, do not preclude the possibility that IL-2 treatment can induce signaling through CD25 as reported (Hashimoto et al., 2022). Thus, IL-2 signaling is likely to be context-dependent. This notion is also highlighted by our data showing that early, in contrast to late, treatment with the IL-2 fusion protein ANV410 abrogated T_PEX_ cell development and impaired the expansion of antigen-specific CD8^+^ T cells.

Recent advancement in our understanding of the central role of IL-2 in the regulation of CD8^+^ T cell expansion and function has reinvigorated the enthusiasm for IL-2-based immunotherapy in infection and cancer. Indeed, IL-2 and IL-2 fusion proteins have emerged as powerful modulators that may promote viral clearance or tumor control. Nevertheless, more work is required to understand the impact of IL-2 in a therapeutic setting on the function of T_PEX_ cells and thus the long-term maintenance of CD8^+^ T cell responses in chronic infection and in cancer. Furthermore, the facts that IL-2 signaling modulated the localization of CD8^+^ T cells and altered the distribution of viral reservoirs which may have implications for people living with HIV or experiencing EBV active infection. Thus, further work is needed to dissect the therapeutic potential of IL-2. Taken together, the development of IL-2-based immunotherapies should not only target the correct receptor subunit but also consider T cell differentiation state and localization.

## Acknowledgments

We acknowledge the Imaging and Cytometry Facility at the John Curtin School of Medical Research, The Australian National University and Flowcores at Monash University and Doherty Institute for technical support. We thank Shane P. Crotty (La Jolla Institute for Immunology) for STAT5b-CA and control vectors, William S Foster and James A. Harker (Imperial College London) for the advice on human studies. We thank ANAVEON for the supply of AN410.

## Author Contributions

Conceptualization: DY and AK

Supervision: DY, AK and ZL

Data curation: YC, PZ, PMG, YAL, JH, YW, FVM, TZ, and JY

Formal analysis: PZ, YC, PMG, YAL, YW and MP

Funding acquisition: DY, AK, PMG and YW

Writing, review and editing: DY, AK, PZ, YC and PMG

## METHODS

### RESOURCE AVAILABILITY

#### Lead contact

Further information and requests for resources and reagents should be directed to and will be fulfilled by the lead contact, Di Yu.

#### Materials availability

This study did not generate new unique reagents.

#### Data and code availability

RNA-seq data were generated from datasets publicly available at GEO. Accession numbers are listed in the Key resources table. qPCR data, microscopy data, and any additional information required to reanalyse the data reported in this paper are available from the lead contact upon request. This paper does not report original code. Scripts and code utilised in this study are available from the lead contact upon request. Information regarding microglia isolation and sequencing can be addressed to Di Yu.

## EXPERIMENTAL MODEL AND STUDY PARTICIPANT DETAILS

### Mice

All mice used in this study were held in SPF (specific pathogen free) animal facilities in The Australian National University, Monash University, University of Melbourne or the Walter and Eliza Hall Institute (WEHI). 6-12 weeks C57BL/6 mice or *Ptprca* (CD45.1) mice were used for *in vivo* and *in vitro* experiments. P14 mice (JAX Tg(TcrLCMV)327Sdz) with the transgenic expression of a TCR specific for LCMV glycoprotein (GP)_33-41_ were used for adoptive transfer experiment, and OT-I mice are with the transgenic expression of a TCR specific for chicken ovalbumin (OVA)_257-264_. *Lck*^Cre^ mice *were crossed with Prdm1^flox/flox^* mice (Kallies et al., 2009). All animal experiment ethics were approved by the designated Animal Ethics Committee of the institutes that were responsible for animal experiments.

### Human data

The details of the prospective, open-label study of low-dose recombinant human IL-2 (rhIL-2) in patients with active SLE was described (He et al., 2016a). Briefly, three cycles of rhIL-2 were administered subcutaneously at a dose of 1 million IU every other day for 2 weeks (a total of 7 doses), followed by a 2-week break. Immunological data in peripheral blood mononuclear cells (PBMCs) were measured at baseline and every 2 weeks thereafter until week 12.

## METHOD DETAILS

### Viral infection

Lymphocytic choriomeningitis virus (LCMV) Docile was propagated using BHK cell line which was cultured with complete DMEM medium. To study the role of IL-2 treatment in chronic LCMV infection, mice were intravenously (i.v.) infected with 1-2×10^6^ PFU (plaque-forming units) of LCMV Docile virus in 100 ul of PBS.

### IL-2 and ANV410 treatment

10,000 to 100,000 I.U. (international units) of human rhIL-2 (recombinant human IL-2^Ser125^, Beijing SL Pharma) were intraperitoneally (i.p.) injected into the mice daily for 5 times from days 10 to 14 post LCMV Docile infection. Mice were sacrificed on day 15 post infection and samples were collected for further examination. ANV410, supplied by Anaveon AG (Basel, Switzerland), is a human IL-2/anti-IL-2 circularly permutated fusion protein which selectively signals through the intermediate affinity IL-2R (IL-2Rβ/γ). For treatment with ANV410, naïve or Docile infected mice were treated with 200 μg/kg body weight three times (every other day in five days).

### Generation of bone marrow chimeric mice

Bone marrow (BM) cells were collected from femurs of 6-12 weeks old mice with indicated genotypes. Single cells were prepared by pipetting in complete RPMI media. CD45.1 recipient mouse were lethally irradiated twice with 4.75 gray (Gy) of gamma radiation 4 hours apart. BLIMP1-deficient BM cells (CD45.2) were mixed with wildtype control BM cells (CD45.1/CD45.2) at 1:1 ratio and intravenously transferred into irradiated CD45.1 recipient mouse. As control, BM cells from genotyped wild type littermates (CD45.2) were also mixed with wildtype control BM (CD45.1/CD45.2) and reconstituted in recipient CD45.1 mice. Mice were housed for 6 to 8 weeks to allow hematopoietic reconstitution before infection with LCMV Docile virus.

### *In vitro* induction of CXCR5^+^ CD8^+^ cells

For induction of CXCR5^+^ CD8^+^ T cells, naïve OT-I cells (CD8^+^Va2^+^CD44^-^CD62L^+^) were purified (10^4^ cells/well) and co-cultured with congenical wildtype splenocytes (10^5^ cells/well) in 96-well plates, in the presence of the SINFEKL peptide (1 µM) with indicated cytokines: TGFβ (10 ng/mL), IL-6 (100 ng/mL) and IL-2 (500 I.U./mL∼30-50ng/mL).

### Retroviral transduction on primary cells

GFP bicistronic retroviral vector containing STAT5b-CA or empty vector as described (Johnston et al., 2012) were used to study the overexpression of STAT5b-CA in primary mouse CD8^+^ T cells. Retroviruses were generated by transfecting the plasmids into a retrovirus-packaging cell line GPE86 using lipofectamine 2000 reagent (Invitrogen). GFP positive GPE86 were cell sorted to isolate stably transfected cells. Stably transfected GPE86 were then cultured in complete DMEM media for 48 hours before the culture supernatant were collected for the transduction of primary cells. Naïve P14 T cells were purified from CD45.2 TCR transgenic P14 mice and stimulated with plate bound anti-CD3 and anti-CD28 for 48 hours. Activated P14 T cells were spinoculated at 800g for 1 hour at 32°C with the retrovirus supernatants packaged by GPE86 cells. Transduced P14 T cells were rested in fresh cRPMI media containing 20 ng/mL of rmIL-7 for 48 hours. About 6,000 GFP^+^ P14 T cells were sorted by flow cytometry and intravenously (i.v.) transferred into CD45.1 recipient mouse. One day after adoptive transfer, mice were i.v. infected with LCMV Docile at 10^6^ PFU. Mice were culled on day 8 post infection for analyses.

### Flow cytometry

Single cells were washed, counted, and stained with Fc-receptor blocking antibodies (clone 2.4G2, 1:100 dilution, BD) for 10 min on ice to block non-specific staining. For surface staining, cells were washed once with PBS containing 2% heat-inactivated fetal bovine serum (FBS, Gibco) and stained in an appropriately diluted antibody solution for at least 30 min at 4 °C. The 7-amino-actinomycin D (7-AAD) (BioLegend) was stained to exclude dead cells. For intracellular staining, Cytofix/Cytoperm (BD) was used as user’s guide. For intranuclear staining, Foxp3/Transcription Factor Staining Buffer Set (eBioscience) was used as user’s guide. To detect LCMV antigen-specific CD8^+^ T cells, APC-conjugated H-2D^b^-GP_33_-tetramer (produced by Biomolecular Resource Facility at The John Curtin School of Medical Research, The Australian National University). To detect LCMV-infected T cells, purified anti-LCMV nucleoprotein (clone VL4) antibody was labelled with Alexa Fluor (AF) 647 using antibody labelling kit (Invitrogen). Splenocytes from LCMV infected mice were first stained with surface antibodies. After permeabilization by the Cytofix/Cytoperm (BD) kit, labelled AF647-VL4 antibody was incubated with the cells for 30 min at 25 °C. Uninfected splenocytes and VL4 unstained splenocytes were used as negative control. Data were collected on a LSR-II or LSRFortessaTM (BD) and analyzed using FlowJo software.

### Immunofluorescence

To visualize LCMV infection and P14 cells in splenic tissues, immunofluorescent staining was performed. The middle section of spleens was cut and submerged immediately in Optimal Cutting Temperature (OCT) compound on dry ice and stored at –80°C. OCT blocks were then sectioned at 5 to 10 μm thickness at –20°C. Sections were blocked with Fc-receptor-blocking antibody (clone 2.4G2, 1:100, BD), followed by staining with primary antibodies. Then sections were washed and stained with secondary antibodies or streptavidin. All steps were performed at 25 °C for 60 min in dark and sections were mounted using Prolong Gold (Invitrogen). Slides were visualized using laser scanning confocal microscopy (Leica SP5) at the John Curtin School of Medical Research. Quantification of infected cells was performed by automatic counting of cells per area of section using Imaris 9.0 software.

### RNA sequencing and bioinformatics analysis

RNA sequencing and bioinformatic analyses to compare the transcriptomes between CXCR5^−^ and CXCR5^+^ P14 cells were described previously (Leong et al., 2016). DESeq2 from Bioconductor software (Love et al., 2014) was used to normalize gene counts in GSE76279 dataset, and the genes were ranked based on –log(adjusted p-value of fold change). The ranked list was introduced to GSEA software (Subramanian et al., 2005) to enrich genes in gene sets of a wide range of databases using a curated version of “msigdb v7.2 symbols”. The normalized counts were also filtered for interleukin receptors to produce a heatmap using pheatmap function in r and to construct a bar chart in GraphPad Prism 9. Multiple unpaired t-tests were used to show the level of significancy.

## QUANTIFICATION AND STATISTICAL ANALYSIS

### Statistical analysis

All results were analyzed by unpaired or paired and two-tailed student t-test for comparison between two groups or one-way ANOVA for calculating univariate data set with more than two groups by GraphPad Prism 8.0 software, unless otherwise specified. Differences were considered to be statistically significant at p < 0.05.

## Supplementary Materials

**Figure S1-S2**

---

**Suppl. Figure 1.**
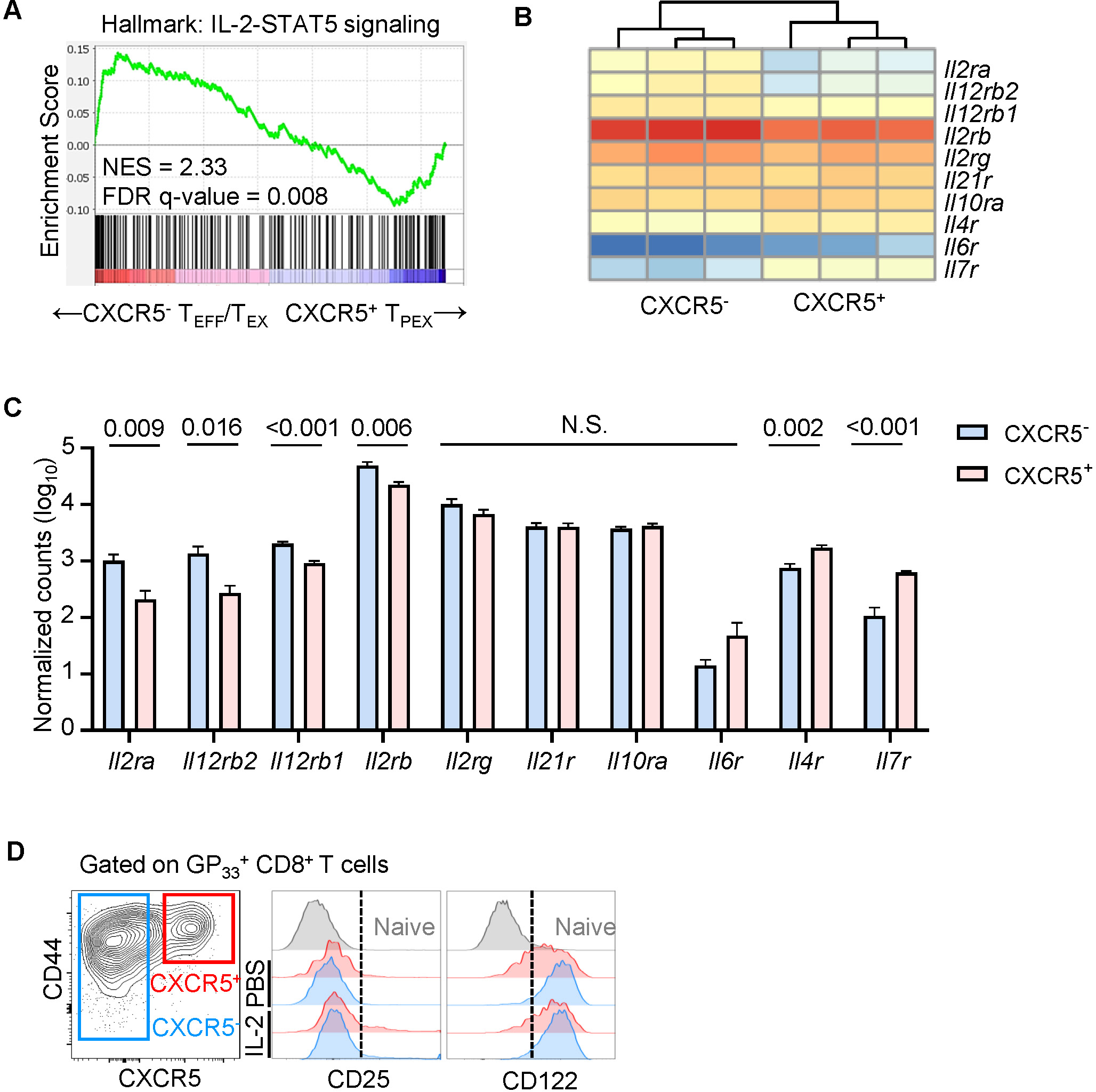
Comparison of IL-2-STAT5 signaling and IL-2 receptors between CXCR5^+^ T_PEX_ cells and CXCR5^-^ T_EX_ cells. (**A-C**) Purified CXCR5^+^ T_PEX_ and CXCR5^−^ T_EX_ P14 cells from host C57BL/6 mice at day 8 post LCMV (DOCILE) infection underwent RNA-seq to generate transcriptomes as described previously (Leong et al., 2016). Gene set enrichment analyses showing the downregulation of IL-2-STAT5 signalling in CXCR5+ T_PEX_ cells (A). Heatmaps (B) and quantification (C) showing the expression of indicated cytokine receptor genes. (**D**) C57BL/6 mice were chronically infected with LCMV Docile and receiving intraperitoneal treatment of IL-2 as in Figure 1A. FACS plots showing the gating of CXCR5^+^ T_PEX_ cells and CXCR5^−^ T_EX_ populations in LCMV-specific GP_33_ tetramer^+^ CD8^+^ T cells (left) and histograms showing the expression of CD122 and CD25 on indicated populations.

**Suppl. Figure 2.**
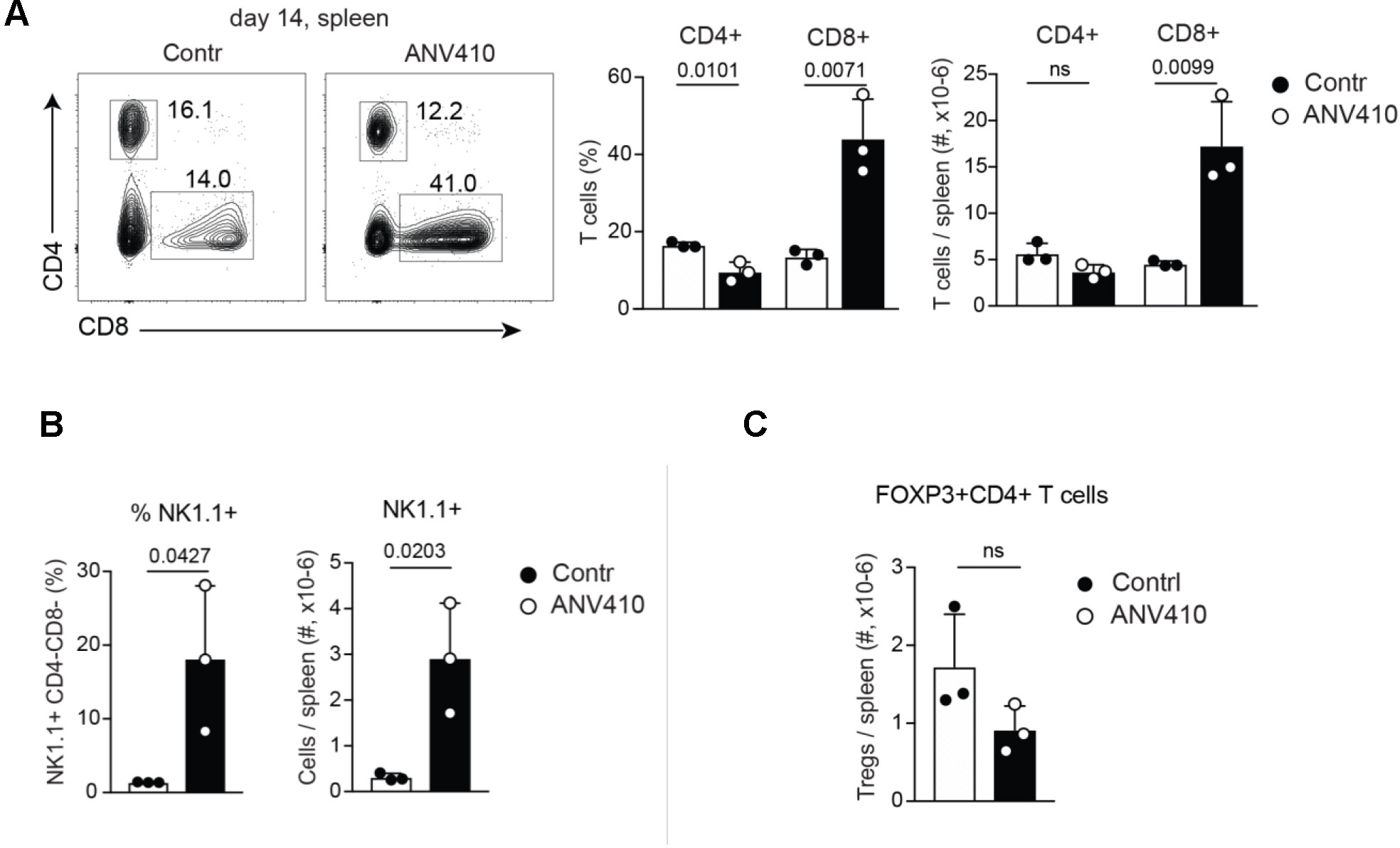
Treatment with a CD122-specific IL-2 fusion protein results in CD8 T cell and NK cell but not T_REG_ cell expansion. C57BL/6 mice were chronically infected with LCMV Docile and receiving intraperitoneal treatment of ANV410 as in Figure 4A (Regimen A). (**A**) FACS plots (left) and quantifications (right) for total CD4^+^ or CD8^+^ T cells. (**B**) Quantification for CD3^-^NK1.1^+^ NK cells. (**C**) Quantification for Foxp3^+^ CD4^+^ T_REG_ cells.

